# Assessment of sexual dimorphism *Desmodus rotundus* (Chiroptera: Phyllostomidae), a rabies reservoir in Latin America

**DOI:** 10.1101/2025.02.17.638589

**Authors:** Analorena Cifuentes-Rincon, Karen D. Sarmiento-Arias, Diego Soler-Tovar, Abelardo Rodríguez-Bolaños, Carlos Bravo-Garcia, Nicolas Reyes-Amaya, Laura Ávila-Vargas, Luis E. Escobar

## Abstract

Morphometric traits of a host have been used recently in disease ecology for a deeper understanding of the connection between phenotype and transmission rates. The common vampire bat, *Desmodus rotundus,* is the main reservoir of rabies in Latin America, one of the most lethal zoonotic diseases in the world. Comprehension of morphological variation in *D. rotundus* is insufficient, contradictory, and inconclusive. Due to this inconsistency, we explored sexual dimorphism in *D. rotundus by combining different measures* to provide new knowledge that can contribute to the control of rabies*. As a result, our study provides evidence of sexual dimorphism in D. rotundus with females generally larger than males in body and wing measurements.* The statistical, as well as the principal components, and clusters analysis, confirm morphological differences between females and males, without observing a complete separation between the sexes, possibly due to environmental factors that change the specimens’ conditions or limitations with access to measures. Understanding the sexual dimorphism of the main transmitter of wild rabies in South America is essential since morphological variations between sexes could influence both the use of the habitat and the dispersion capacity of the bats, which are relevant to the epidemiology of rabies. Because our data originated primarily from lowland areas, we are unable to rule out the effects of elevation on sex dimorphism. Given the importance of rabies for global public health, investigating the morphological and behavioral aspects of rabies reservoirs could help us better understand how ecological aspects influence disease spread.

## Introduction

Morphometric analysis is rutinary applied to comparative biology studies to better understand ecological and evolutionary trajectories. Phenotypic traits of a host have been used recently in disease ecology to better understand linkages between morphological characteristics and transmission rates [1]. For instance, current literature suggests that the number of pathogens increases with the body size of the host and that larger mammals have a stronger immune response against infection than smaller mammals [2–6]. Body size patterns do not occur randomly, some authors have demonstrated reductions in the body mass of bats across progressive pathogen exposure [7]. As such, morphological variation in hosts could inform how host individuals participate in pathogen transmission, maintenance, and spread. Research on morphological variation at the species level in the context of disease ecology, however, remains poorly explored.

*Desmodus rotundus* (E. Geoffroy, 1810), is one of the most extensively studied bat species in the context of disease ecology [8]. Numerous pathogenic viruses, some of which are highly dangerous, have been identified in *D. rotundus* across the Neotropics [9–11]. *Desmodus rotundus* is the main source of transmission of rabies in Latin America [5,6]. Rabies is considered one of the most fatal zoonotic diseases globally [12, 13]. Understanding of morphological variation in *D. rotundus* regarding sex dimorphism is incomplete and heterogeneous. For example, some authors have reported females being larger than males in terms of sexual size dimorphism [14–17]. Other authors found males larger than females in terms of sexual selection, as well as indicating that forearm length in bats could be greatly influenced by phylogenetic relationships [18]. Additionally, there are reports of no significant differences between *D. rotundus* females and males, with suggestions that morphological variation may be influenced by food availability [19]. The understanding of sexual dimorphism in *D. rotundus*, as a form of morphological variation, remains inconclusive.

Due to the discrepancies in the literature and the importance of this species in rabies transmission, we explored sexual dimorphism in *D. rotundus* to provide valuable knowledge for improving future rabies control efforts. We aimed to address the question: Is there sexual dimorphism in the common vampire bat, *Desmodus rotundus*? We examined newly acquired specimens from 2022 and 2023, along with historical samples from museum specimens collected over a century (1921 to 2023) available in 11 different collections. Our study focused on *D. rotundus* populations along Colombia, a country offering large elevational and climatic gradients.

## Methods

We collected morphometric data from 60 *D. rotundus* individuals (43 males and 17 females) at five field sites from 2022 to 2023 across Colombia (Fig. 1A). Bat capture and sampling procedures were conducted in accordance with ethical guidelines for wildlife research. All methods were approved by Virginia Tech IACUC-21-138, and Universidad de La Salle – Colombia. Permit for Collection of Specimens of Wild Species, 1473. Sampling was carried out with the authorization of the Ministry of Environment of Colombia and complied with national and international regulations for the ethical treatment of wild animals.

**Figure 1.**
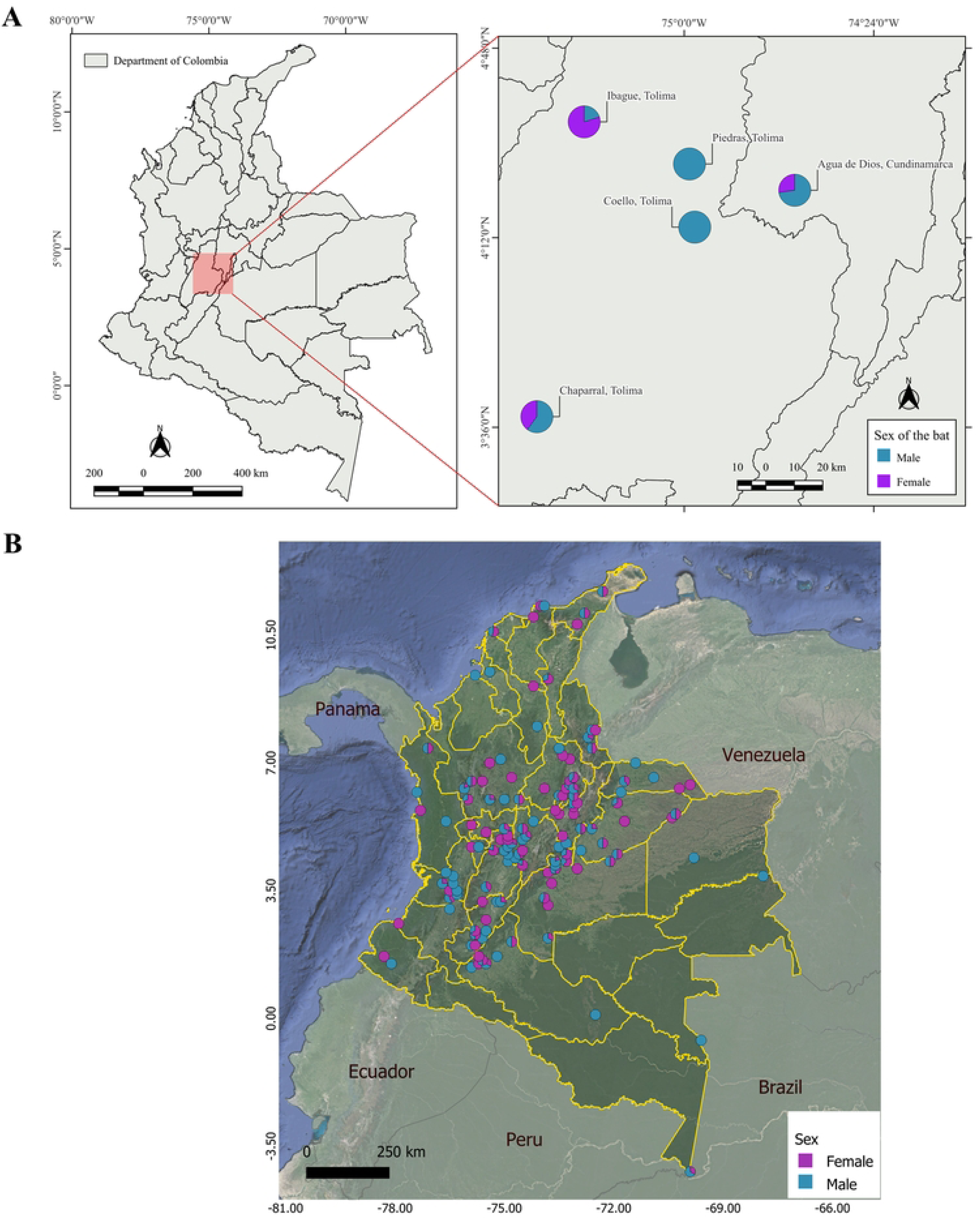
Geographical representation of the records of *Desmodus rotundus* in Colombia, represented by sex. **A.** Geographic distribution of 60 *D. rotundus* individuals from which six morphological traits were collected (i.e., weight, head length, body length, tibia length, ear length, and forearm length.) in the departments of Tolima and Cundinamarca. **B.** Distribution of 490 *D. rotundus* museum specimens from which forearm measurements were collected for this study. Violet: female. Blue: male.

Bats were captured using mist nets and handled following established protocols to minimize stress and ensure their welfare. After sample collection, individuals were released at the site of capture. No endangered or protected species were included in this study.

To assess sexual dimorphism we selected weight, head length, body length, tibia length, ear length, and forearm length as focal metrics (Table 1). These measurements are standard in bat taxonomy and provide a robust source for exploring morphological variation. Each measurement was chosen based on its biological relevance, accuracy in differentiating sexes, and frequent use in chiropteran studies [20].

**Table 1.**
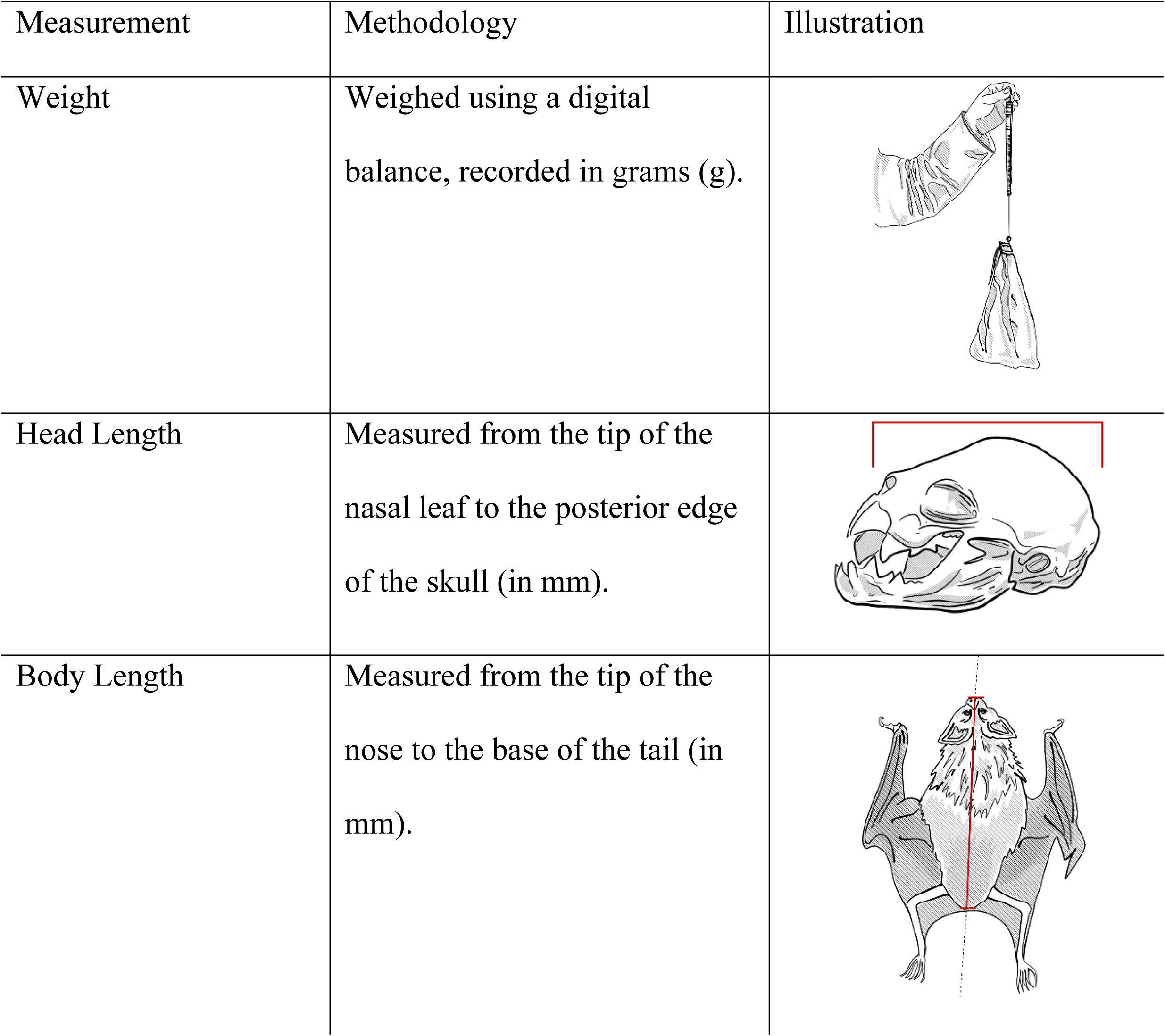
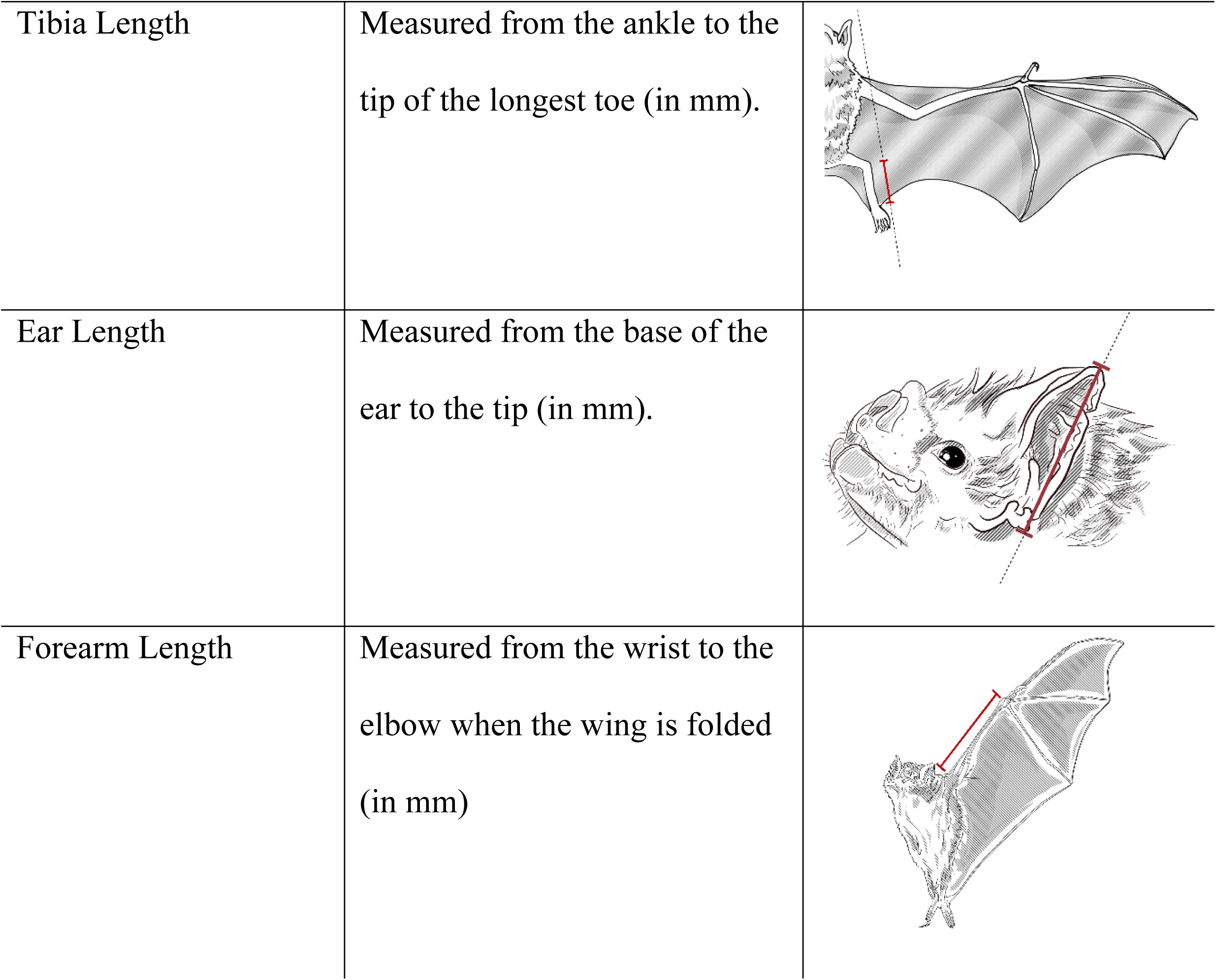
Morphological measurements of Desmodus rotundus.

| Measurement | Methodology | Illustration |
| --- | --- | --- |
| Weight | Weighed using a digital balance, recorded in grams (g). |  |
| Head Length | Measured from the tip of the nasal leaf to the posterior edge of the skull (in mm). |  |
| Body Length | Measured from the tip of the nose to the base of the tail (in mm). |  |
| Tibia Length | Measured from the ankle to the tip of the longest toe (in mm). |  |
| Ear Length | Measured from the base of the ear to the tip (in mm). |  |
| Forearm Length | Measured from the wrist to the elbow when the wing is folded (in mm) |  |

This table presents the various morphological traits measured in *D. rotundus*, detailing the methodology employed for each measurement. Illustrations are included, with each measurement highlighted in red to visually demonstrate the precise locations of the measurements.

Bats were collected using three capture methods (i.e., conventional mist nets, harp trap, and cone trap) [21,22] and placed in cloth bags. A spring balance was used to weigh each bat and the breeding stage was reviewed by selecting pre-weaned (bat during the early lactation period), juvenile, subadult, or adult. Adults were classified based on their reproductive condition (females were classified as active or inactive, lactating or pregnant, and males as having inguinal or descended testicles).

Subsequently, individuals were measured using a Uline digital caliper, with an accuracy of 0.0005mm. Individuals included in our analyses were only those belonging to the adult and semi-adult categories, excluding specimens classified as sexually immature or pregnant to avoid generating noise in the analysis. Specimens were confirmed to be the species *D. rotundus* using broadly used taxonomic keys [23–25].

We performed a statistical analysis of quantitative morphological data using the prcomp function, stats package in R [26]. Analyses included principal component analysis (PCA) and multivariate clustering using the Euclidean distance parameter to measure the similarity or difference between *D. rotundus* in Colombia through continuous variables. We used a PCA to rescale data while preserving essential information, simplifying analysis from fewer dimensions. This approach facilitated visualization in two axes that retain most information from the original data and eliminate redundancy between variables to visualize a separation by sex.

Hierarchical clustering using hclust function in R allowed us to group individuals into sets with similar morphological values. Clusters were visualized using a dendrogram denoting morphological relationships between sexes of *D. rotundus* individuals using the dendextend package in R [26], where we use sex as a predictor variable and morphometry as a response variable. Finally, the correlation between ear and forearm, forearm and weight, and body and weight measurements were measured to corroborate the influence of the variables.

In bats, forearm length (the segment of the wing extending from the elbow to the wrist) is the most reliable and widely used measurement to assess size [27,28]. This measurement is diagnostically valuable due to its involvement in aerodynamics, ecological factors, geographic variation, and sexual dimorphism [29]. For this study, we acquired additional forearm length data from 490 specimens (210 females and 280 males) in museum collections (Fig. 1B). Forearm data were employed alone to estimate sexual dimorphism from these individuals across Colombia (Fig. 1B).

We visited 11 museums from Colombia, including Museum Collection of Universidad de La Salle -MLS-mam [30], Zoological Collections Museum of Natural History Universidad de los Andes -ANDES-M [31], Mammal Collection “Alberto Cadena García” Universidad Nacional de Colombia -ICN [32], Alexander von Humboldt Institute Collection -IAvH [33], Museum of Natural Sciences Universidad de La Salle -CSJ-m [34], Museum of Natural History Universidad de la Amazonía -UAM, Natural History Museum Universidad de Los Llanos -MHNU-M [35], Mastozoology Collection of the Natural History Museum Universidad Industrial de Santander -MHN-UIS, Zoological Collection Universidad del Tolima -CZUT-M, Mammal Collection of the Museum of Natural History Universidad de Caldas -MHN-Uca [36], and Mammal Collection, Natural History Museum Universidad Distrital Francisco José de Caldas (MHNUD-M) [37]. We measured the forearm length of specimens of *D. rotundus* from museum collections between October 2022 and April 2023. All forearm length measurements were taken using a 0-150mm 6″ PRD electronic digital caliper, with an accuracy of 0.01mm.

Using R statistic software version 4.1.2 [26] and RSTUDIO (ver. 2022.02.3), we used a student t-test [38] in the BayesFactor package [39]. We also performed a boxplot using the geom_boxplot function in the ggplot2 package [40], and hierarchical clustering using the distance matrix hclust, the hierarchical grouping was converted into a phylogenetic tree through the ape package [41]. Finally, the hierarchical tree was edited in the Interactive Tree Of Life (ITOL V6) online program [42

## Results

Results suggest sexual dimorphism between males and females of *D. rotundus*, the females being larger than the males. This is supported by two statistical analyses: one incorporating a series of morphological metrics and another focusing exclusively on forearm length. The principal components analysis conducted on a sample of 60 individuals using six morphological measurements (body weight, head length, body length, tibia length, ear length, and forearm length), revealed significant variability in body measurements attributable to sex (χ²=866.26, df=20, p=1.18×10⁻¹⁷⁰). This indicates a significant effect of sex on body morphology. Few outliers were found in individuals from one study site (i.e., Agua de Dios, Cundinamarca). These outliers showed morphological differences from the bulk of samples (Fig. 2A, B).

**Figure 2.**
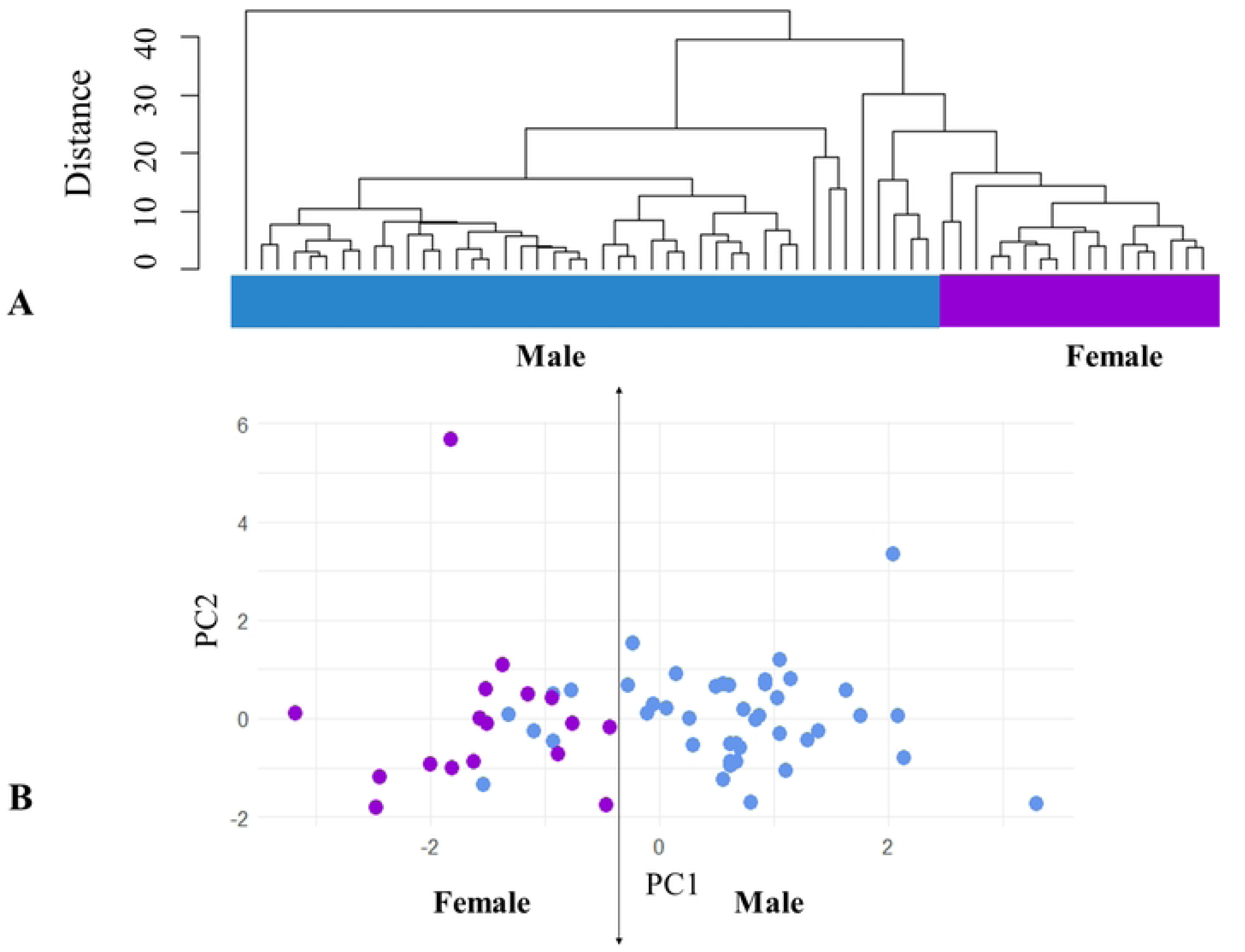
Cluster analyses of bat morphology. **A.** The distance tree by cluster analysis of *D. Rotundus* according to head, body, leg, ear, forearm, and weight derived from 60 individuals. Violet: females. Blue: males. Note how 2 clades tend to form, however, it does not show enough segregation between males and females. **B.** Principal components graph generated using the biometric measurements of *D. Rotundus* individuals showing the distribution of 60 individuals across weight, head length, body length, tibia length, ear length, and forearm length. Violet: female. Blue: male. Isotropy in principal component space suggests that morphological differences between males and females are not limited to a single measurement, but affect multiple aspects of morphology equally.

In the second analysis, which focused on forearm length measurements only in a larger sample of 490 museum specimens, also significant differences were found between males and females (t=-12.677, df=366.8, p=2.2×10⁻¹⁶) with females larger than males. Furthermore, hierarchical clustering based on this trait also demonstrated a separation between males and females (Fig. 3A, B).

**Figure 3.**
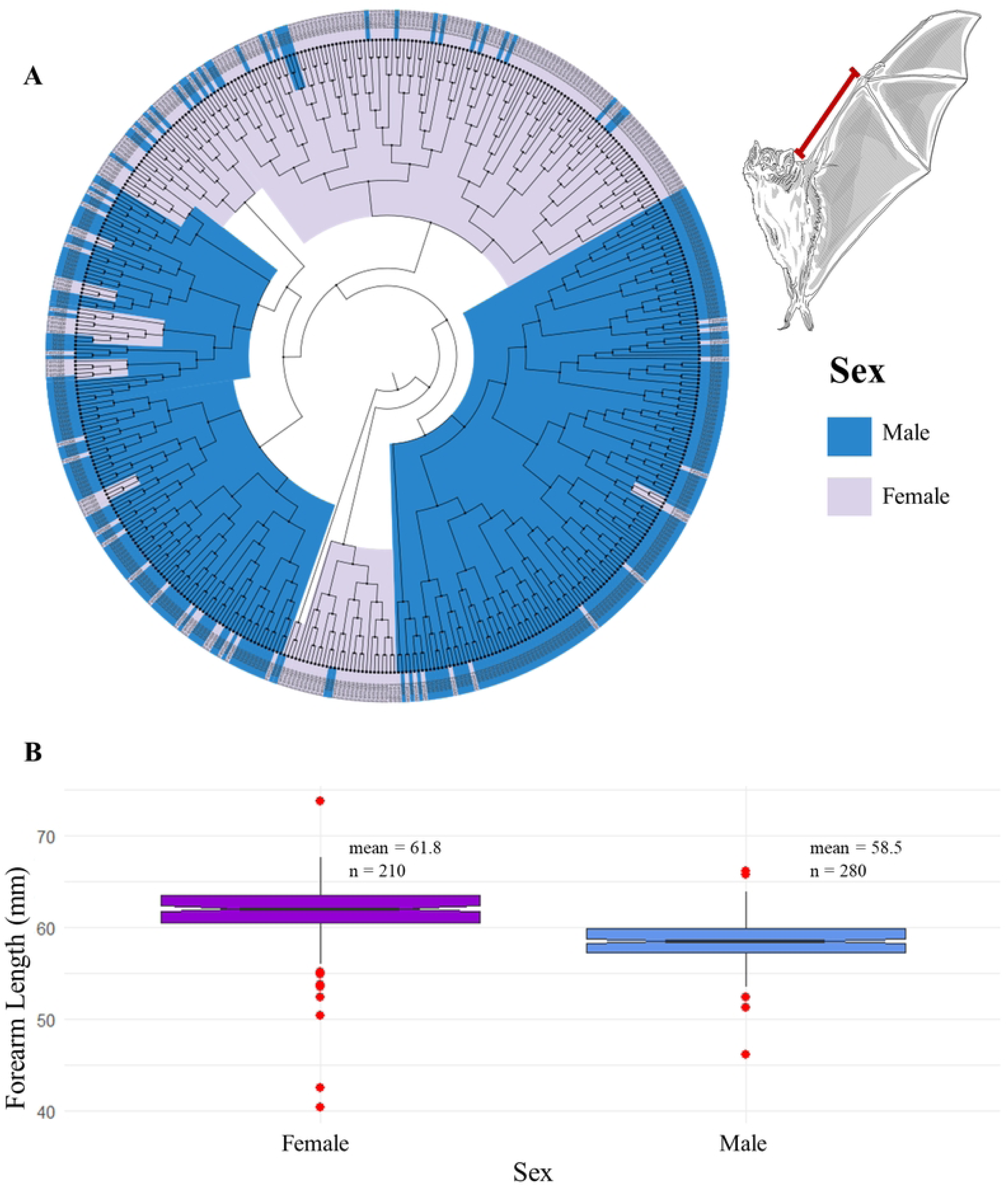
Comparison of forearm length in *Desmodus rotundus* by sex. **A.** Distance tree by cluster analysis of *D. Rotundus* according to forearm length from 490 museum specimens. Violet: females. Blue: males. Note that females and males tended to cluster in two groups. **B.** Boxplot illustrating sexual dimorphism in forearm length, displays the median (central line) higher on the female box plot compared to the male. Females generally show longer forearm lengths than males, the interquartile range is similar, slightly larger for females, signifying a wider distribution of forearm lengths within the female population; outliers (red points), appear beyond the whiskers, showing a few individuals with significantly shorter or longer forearms compared to the majority within their sex group.

## Discussion

Sexual dimorphism was found in both analyses (i.e., one using various body measurements in 60 individuals and the forearm analysis using 490 individuals). Sexual dimorphism was most pronounced when measuring forearm length alone, with females consistently exhibiting larger measurements than males. Outlier specimens falling in the range expected for the opposite group can be due to aspects such as physical condition, age, and environmental factors [43,44]. Measurement errors could also explain incomplete sexual dimorphism [45,46]. Geometric morphometry assessments based on analysis of digital imaging technology could refine studies of sexual dimorphism and could address the limitations of our study for more accurate measurements [47,48].

Our statistical analyses reveal significant differences in body size between females and males in *D. rotundus*, corroborating previous studies that reported females as generally larger [14–17]. Specifically, forearm measurements from 490 museum specimens, demonstrated a highly significant distinction between sexes (t=-12.677, df=366.8, p=2.2×10⁻¹⁶). Furthermore, the principal component analysis of all morphological measurements of the body indicated that variability in body size is strongly influenced by sex (χ²=866.26, df=20, p=1.18×10⁻¹⁷⁰). These findings support the sexual dimorphism in *D. rotundus* and establish forearm length as a dependable trait for distinguishing between male and female.

This pattern may be related to the big mother theory, where evolutionary selection is hypothesized to favor large females assumed to have more physiological resources for the offspring [49]. For instance, female *D. rotundus* bats exhibit feeding site displacement over males and take advantage of bites from other bats by directly flying over prey and conserving energy [50,51]. A larger body size would facilitate competition within species, as revealed by *D. rotundus* displacement [17,50].

Stevens et al. [49] studied sexual dimorphism using wing and body size in the species frugivorous bat *Artibeus lituratus* and found sexual differentiation. *Artibeus lituratus* revealed a pattern of larger females when the size of wings was considered, but no difference when a phenotypic integration was used (i.e., multiple body measurements). Larger wings of females, in the absence of overall sexual dimorphism, could be due to greater aerodynamic requirements in response to the increased weight of females during reproduction [52], which has also been observed in birds [53]. Flying mammals differ from birds in the duration, speed, and direction of flight, all of which dictate wing differences from aerodynamic pressures [54]. Similarly, O’Mara et al. [55] found sexual dimorphism in the insectivorous bat *Nyctalus 13lock13t*, where larger females were observed in terms of wing size and body mass. In the case of *N. 13lock13t*, sexual dimorphism could be linked to differences in behavior, since only females migrate. Morphological differences have been attributed to size, but not wing shape suggesting that sexual dimorphism is likely due to the reproductive advantage of larger female size [56,57].

Females of *Myotis bechsteinii* have been observed to exhibit fluctuating asymmetry (e.g., random, non-directional variations in bilateral morphological traits) and directional asymmetry (i.e., consistent differences where one side is larger than the other on most individuals in the population) in their forearms, without negative effects on their reproductive success [58]. Wing asymmetry could be associated with environmental stress or anthropogenic disturbances [59], with sex likely influencing patterns of asymmetry [60]

Ueti et al. [60] observed that in *D. rotundus* morphometric asymmetry varied with sex, females exhibiting fluctuating asymmetry and males exhibiting both directional asymmetry and antisymmetry (i.e., the larger side varying randomly between individuals). Our results show that females of *D. rotundus* are larger than males without assessment of asymmetrical variability, as all forearm measurements were taken exclusively from the right forearm. Nevertheless, we can see from these findings the importance of size differences between sexes, which may be associated with different reproductive and ecological pressures on *D. rotundus* females vs. males.

Because most individuals (65%) for this study were collected at low elevations (i.e., <1000 m.a.s.l), we were unable to robustly assess the effect of elevation on sexual dimorphism in *D. rotundus*. Nevertheless, other studies have revealed that birds and bats display differential sex-related thermal preferences where females prefer higher temperatures than males [61]. Thermal preferences in bats have been proposed as a driver of geographical segregation of males and females in the insectivorous bat *Rhinopoma microphyllum* [62]. During summer, *R. microphyllum* males tend to feed at colder temperatures, whereas females prefer to forage in the lowlands [63].

Accounting for elevational gradients would improve our understanding of thermal adaptation that contributes to morphological differences, geographic segregation, and demographic parameters in *D. rotundus.* Some species reveal a correlation between morphology and habitat use [64]. Temperature and humidity preferences could be linked to the physiological adaptation of bats to establish maternity colonies in warmer roosts [65]. This information would be critical to understanding how future climate changes would impact the ecology of *D. rotundus* and, in turn, the epidemiology of rabies.

Rabies is a zoonotic disease produced by rabies virus, which is found in a variety of animals, primarily in bats and other species from the order Carnivora [66]. Bat-borne rabies infections occur in domestic animals and humans where infection is usually fatal [67,68]. Rabies causes 60,000 human deaths annually, which means one death every 10 minutes approximately, especially in Asia and Africa [69]. Accordingly, comprehending the sexual dimorphism in a disease ecology context of a rabies reservoir could help improve our awareness of rabies epidemiology.

Rabies host distribution could be influenced by morphological and behavioral features. Morphological differences between females and males can impact their movement patterns within the environment [70]. Movement variations can influence the transmission dynamics of rabies, while host ecology also plays a crucial role in rabies spread. Body size is important in disease ecology considering that larger animals are considered more susceptible to transmit zoonotic viruses [71–73].

Some authors have suggested male-biased gene movement and female philopatry in temperate bats [74,75], including *D. rotundus* [76,77]. The fact that females are larger than males, as suggested in this study, could influence the social structure of *D. rotundus* and, therefore, the population’s genetic structure and rabies dissemination [78]. Thus, sex-biased surveillance targeting more samples from females could help improve rabies detection and monitoring. This hypothesis, however, needs to be tested in field conditions.

We observed female-based sexual dimorphism using measurements related to body size, such as weight, body length, and forearm length, as measurements related to echolocation such as ear length, and those related to terrestrial locomotion, such as tibia length. Strong terrestrial locomotion allows *D. rotundus* to approach their prey carefully and quickly escape with flight-initiating potent jumps from any threat while feeding [79]. This ability is crucial for *D. rotundus*, as their feeding habits involve consuming animals heavier than them, while also exposing them to predators [80]. Because females are generally larger, their increased body size, especially in forearm length, may confer greater mobility and flying ability.

Additional measurements related to flight, terrestrial locomotion, and feeding behavior, such as digit length, wing loading, aspect ratio, wingspan, paw length (with and without claws), pollex length (with and without claws), tragus length, and nasal leaf size, could have provided deeper insights into the morphological differences between female and male *D. rotundus*. These traits might offer a clearer understanding of how these differences impact their role as vectors of rabies virus. Unluckily, these measurements were not available for incorporation into our analysis.

## Conclusion

Our study on *Desmodus rotundus* supports sexual dimorphism, with females generally larger than males, as observed through forearm length and body size metrics. Although the statistical analyses suggest morphological differences between sexes, a complete separation was not evident. The morphological differences that we propose in this study could explain differences in the use of the habitat between sexes, and consequently, impact the dynamics of rabies. For instance, differences in forearm could affect prey or habitat preferences, and influence the likelihood of contact with other species, potentially altering the spread of the rabies. Incorporating morphological and behavioral factors into ecological and epidemiological models could provide a deeper understanding of how physical traits influence rabies transmission dynamics. Given the global significance of rabies for public health, further investigation into the link between morphology and disease spread in rabies reservoirs could yield important insights for managing and mitigating the disease.

## Acknowledgments

We extend our gratitude to the present and past directors, curators, and managers of natural history collections in the museums in Colombia for facilitating access to specimens. We thank Julieth Cardenas, Gelys Mestre, Fernando Sarmiento, Andrew Jackson, Sebastián García, Catalina Cárdenas, Catherine Mora, Hugo Fernando López, Sandra Galeano, Andres Lozano, Andrea Bustamante, Alexander Velasquez, Oscar Marin, Elizabeth Aya, Francisco Sanchez, Victor Serrano, Giovany Guevara, Gladys Reinoso, Leidy Ramírez, Leidy García, Hector Ramirez, and Catalina Torres for their contributions. We thank Paige Van de Vuurst, Sharif Islam, Quan Dong, Mariana Castañeda, and Daniel Rendón. The authors thank local guides, including Jose Trujillo and Diego Ceballos, and community members for assisting us during fieldwork. This project was supported by National Science Foundation CAREER (2235295) and HEGS (2116748) awards, Virginia Tech DA PPP CeZAP, and ICTAS grants. Research reported in this publication was supported by the National Institute of Allergy and Infectious Diseases of the National Institutes of Health under Award Number K01AI168452. The content is solely the responsibility of the authors and does not necessarily represent the official views of the National Institutes of Health.

## Author contribution

ACR: Data curation, Formal analysis, Investigation, Methodology, Visualization, Writing – Original Draft Preparation, Writing – review & editing. KDSA: Data curation, Validation. DST: Project Administration. ARB and NRA: Resources, CBG: Data curation, Resources. LAV: Data curation. LEE: Funding acquisition, Project Administration, Conceptualization, Methodology, Supervision, Writing – review & editing.

S1. Alternative Language Abstract (Spanish)

S2. Dataset: Morphometric Measurements of *Desmodus rotundus* Specimens

All authors agree to its submission and the Corresponding author has been authorized by co-authors. This Article has not been published before and is not concurrently being considered for publication elsewhere (including another editor at PLoS One). This Article does not violate any copyright or other personal proprietary right of any person or entity and it contains no abusive, defamatory, obscene, or fraudulent statements, nor any other statements that are unlawful in any way.

